# Three-photon imaging of synthetic dyes in deep layers of the neocortex

**DOI:** 10.1101/2020.06.04.135442

**Authors:** Chao J. Liu, Arani Roy, Anthony A. Simons, Deano M. Farinella, Prakash Kara

## Abstract

Multiphoton microscopy has emerged as the primary imaging tool for studying the structural and functional dynamics of neural circuits in brain tissue, which is highly scattering to light. Recently, three-photon microscopy has enabled high-resolution fluorescence imaging of neurons in deeper brain areas that lie beyond the reach of conventional two-photon microscopy, which is typically limited to ~450 μm. Three-photon imaging of neuronal calcium signals, through the genetically-encoded calcium indicator GCaMP6, has been used to successfully record neuronal activity in deeper neocortical layers and parts of the hippocampus. Bulk-loading cells in deeper cortical layers with synthetic calcium indicators could provide an alternative strategy for labelling that obviates dependence on viral tropism and promoter penetration. Here we report a strategy for visualized injection of a calcium dye, Oregon Green BAPTA-1 AM (OGB-1 AM), at 500–600 μm below the surface of the mouse visual cortex in vivo. We demonstrate successful OGB-1 AM loading of cells in cortical layers 5–6 and subsequent three-photon imaging of orientation- and direction-selective visual responses from these cells.

## 1. Introduction

The advent of two-photon microscopy^1^ has revolutionized our ability to record the structure and function of various parts of the neocortex at high resolution *in vivo*. In particular, two-photon imaging of cellular calcium signals has allowed monitoring of neuronal activity from a population of neurons while maintaining single-cell resolution^2–3^. Such studies have proven extremely valuable in unravelling the functional organization of the sensory cortices in a wide variety of mammals such as rodents^4–5^, cats^6–8^, ferrets^9–10^ and macaques^11–12^. However, due to the limitation of two-photon imaging of neural activity in the brain to within a depth of 450 μm^13–15^, these experiments have been largely restricted to neocortical layers 2/3. Therefore, the functional micro-organization of neural networks in cortical layers 4, 5 and 6, or in any area below the neocortex remains elusive.

Recently, three-photon imaging has emerged as a powerful technology enabling high-resolution optical imaging in deeper brain areas^16^. With three-photon excitation, a fluorophore transitions to the excited state via the simultaneous absorption of three photons each containing one-third the energy required for the transition, followed by conventional fluorescence emission^17–18^. Therefore, while two-photon excitation uses excitation light with twice the wavelength of one-photon excitation, three-photon imaging uses excitation light with three times that wavelength. The use of longer-wavelength excitation light results in reduced loss of photons due to scattering in the brain and consequently causes less bulk-heating of the brain. In addition, due to the dependence of three photon excitation on the third power of laser intensity, the effective excitation is localized to an extremely thin focal volume, thereby reducing out-of-focus fluorescence from superficial depths and improving signal-to-background ratio (SBR). Utilizing this technique, neuronal calcium responses have been recorded through the entire depth of the visual cortex^19–20^ and from the superficial neurons of the hippocampus^21^ in mice.

To date, all published studies of three-photon imaging of neuronal activity have used the genetically-encoded calcium indicator (GECI) GCaMP6, either in a transgenic mouse^20, 22^ or introduced into the cortical tissue via transfection of AAV-based viral vectors^19, 21^. While the viral approach circumvents the dependence on transgenic availability in a given species, successful labelling of neurons in the targeted brain area depends on two factors: 1. tropism of the specific viral serotype, and 2. the penetration of the promoters being used to drive the expression of the GECI. Failure in either one of these 2 factors could lead to compromised labelling in a particular cell type. For example, transfection with GCaMP constructs packaged into most AAV serotypes and using the synapsin promoter result in a small fraction of neurons in layer 4 being labeled across a variety of species (see Discussion). Labelling in other cortical layers is strong, except that the fraction of inhibitory neurons expressing GCaMP6 is small^23^. An alternative strategy to introduce calcium indicators into a desired brain area without dependence on transfection/expression systems is offered by bulk-loaded synthetic calcium indicators such as Oregon Green 488 BAPTA-1 AM (OGB-1 AM). Such dyes have been successfully used in the past to label and record activity from layers 2/3 neurons in the neocortex through two-photon imaging. In our experience, proper standardization of the dye preparation and injection method can achieve high success rate with dense labelling of cortical networks using these dyes^4, 7, 24^. Therefore, to extend the use of bulk-loaded calcium indicators to deep-tissue three-photon imaging of calcium signals, we standardized a method to inject OGB-1 AM into the deeper cortical layers in the mouse visual cortex and optically recorded visual responses using three-photon calcium imaging. Here we describe the optimization of the various steps involved in the process and demonstrate three-photon imaging of visual responses using OGB-1 AM.

## 2. Methods

### 2.1 Multiphoton imaging setup

Imaging was performed with a customized microscope from Bruker, with two separate excitation sources for two- and three-photon imaging (see Figure 1). An optical parametric oscillator (Insight X3, Spectra Physics) served as the two-photon excitation source. A noncollinear optical parametric amplifier (NOPA, Spectra Physics) running at 1 MHz repetition rate pumped by a 70 W laser (Spirit 1030-70, Spectra Physics) was used as the three-photon excitation source. All structural and functional imaging in the mouse visual cortex and collection of fluorophore brightness spectra (see below) were performed using a 25× objective lens (XLPLN25XWMP2, NA 1.05, Olympus). In addition, a 40× objective lens (LUMPLFLN 40XW, NA 0.8, Olympus) was used during the visualized injection of fluorescent dyes into layers 5–6 of the mouse visual cortex *in vivo* (see below). The emitted photons were separated into green (OGB-1 AM or third harmonic generation) and red (Texas Red Dextran) channels by passing through first a dichroic beam splitter (T565lpxr, Chroma) and then through barrier filters (green: 525 ± 25 nm band pass, Chroma; red: 675 ± 75 nm band pass, Chroma) before being collected by two photomultiplier tube (PMT) detectors (H10770PB-40 SEL, Hamamatsu). For 1300 nm excitation beam from the NOPA, we compensated the group velocity dispersion of the system by placing a two-prism compressor in the optical path before the microscope. No external compensation mechanism was used for 1600 nm excitation beam. The final pulse widths under the Olympus 25× objective, as measured by an autocorrelator (Carpe, APE) using secant-squared fitting, were found to be 65 fs and 58 fs for 1300 and 1600 nm beams, respectively.

**Figure 1.**
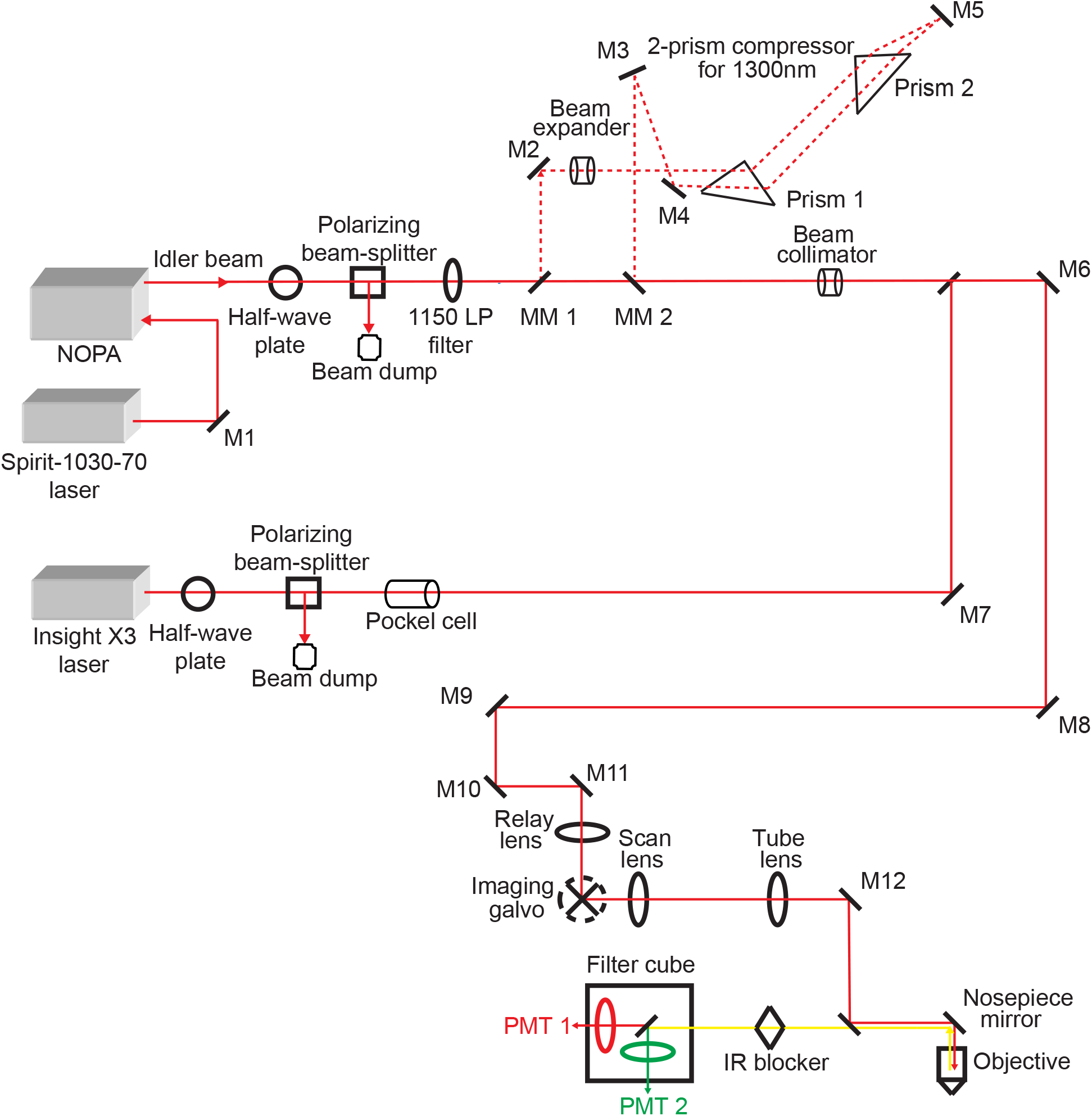
Schematic of the system setup. The 1150 LP filter is used to block the residual visible light from the NOPA. The 2-prism compressor is used to obtain short pulses out of the objective lens for 1300 nm excitation. NOPA, noncollinear optical amplifier; LP, long pass; M, mirror; MM, motorized mirror; IR, infrared; PMT, photomultiplier tube.

### 2.2 Measuring two- and three-photon fluorescence brightness spectra

Selected red and green fluorophores of interest were prepared as aqueous solutions: Alexa Fluor 633 hydrazide 87 μM (Invitrogen); Sulforhodamine 101 7.6 μM (Sigma Aldrich); Texas Red Dextran 70 kDa 87 μM (Invitrogen); mCherry 91 μM (OriGene); mRaspberry 75 μM (OriGene); tdTomato 4 μM (OriGene); Oregon Green 488 BAPTA-1 hexapotassium salt 45 μM (Invitrogen); Fluorescein Dextran 2000 kDa 12.5 μM (Sigma Aldrich). Each fluorophore solution was loaded into a glass micropipette and imaged under the objective lens by varying excitation wavelengths at 10 nm steps and maintaining the same average power on the sample at each wavelength. The fluorescence brightness was calculated by averaging the pixels within a region of interest (ROI) drawn within the pipette and subtracting from it the average brightness within an ROI of the same size drawn outside the pipette. For two-photon spectra of all fluorophores, a wavelength range of 800 to 1200 nm was scanned. For three-photon spectra, 1250 to 1350 nm was scanned for green fluorophores and 1500–1800 nm was scanned for red fluorophores. These spectral measurements were repeated at least twice for each fluorophore.

### 2.3 Animals and surgery

All animal procedures were approved by the Institutional Animal Care and Use Committee of the University of Minnesota. OGB-1 AM injections in deep cortical layers were performed in male C57BL/6J mice (n = 5, postnatal days 71–79). Following dye injections, structural imaging was performed in two of five mice, whereas the other three mice were used for functional imaging of visual responses. For the two mice used for structural imaging, anesthesia was induced by an intraperitoneal injection of urethane (1.5 g kg^−1^), supplemented by injections of ketamine (10–30 mg kg^−1^) and xylazine (1–3 mg kg^−1^), and maintained by one subsequent injection of urethane (0.25 g kg^−1^) during imaging. In the mice used for functional imaging, anesthesia was induced by an intraperitoneal injection of fentanyl citrate (0.04–0.06 mg kg^−1^), midazolam (4–6 mg kg^−1^) and dexmedetomidine (0.2–0.25 mg kg^−1^) cocktail. During functional imaging, a lower dosage of the anesthetic mixture (0.006–0.02 mg kg^−1^ h^−1^ fentanyl citrate, 0.6–2.0 mg kg^−1^ h^−1^ midazolam and 0.03–0.1 mg kg^−1^ h^−1^ dexmedetomidine) was administered through a i.p. catheter connected to a syringe pump. In all mice, the heart rate, temperature and respiration rate were continuously monitored throughout the duration of the experiment. Small craniotomies (2–3 mm in diameter) were made over the primary visual cortex (V1) centered approximately 2.5 mm lateral to the lambda suture and 1–1.5 mm anterior to the transverse sinus. In mice used for structural imaging, Texas Red Dextran (2.5%, 1 μL/g) was injected retro-orbitally to visualize the vasculature.

### 2.4 OGB-1 AM dye injection

#### 2.4.1 Pipette and OGB-1 AM dye preparation

Pipettes for OGB injections were pulled from borosilicate glass capillaries (1B150F-4, outer diameter 1.5 mm, inner diameter 0.84 mm, World Precision Instruments) using a micropipette puller (P-97, Sutter Instrument). The pipettes had a tip diameter of 2–3 μm and a gradual taper such that the diameter at 1.4 mm behind the pipette tip measured ~110 μm. The OGB-1 AM dye mixture was prepared and loaded into the pipettes following procedures that have been described previously^24^. Because the AM ester is transparent until it enters cells, Alexa Fluor 633 hydrazide (Alexa 633) was included in the pipette to visualize the pipette tip during the approach to cortical layers 5–6 and through the ejection of the AM dye (see below).

#### 2.4.2 Determining the entry point of the pipette into the brain

The mouse was positioned on a stage under the microscope and the head was tilted ~15° with respect to the imaging plane to provide full clearance for the pipette to approach the cortical surface without bumping the objective lens. The location for the pipette’s entry into the brain was first determined by visualizing the cortical surface through a 40× objective lens (3.3 mm working distance) under bright-field illumination. The angle for the pipette’s approach was set to 30° with respect to the imaging plane (thus, ~45° relative to the brain surface due to the tilting of the head). During travel to the injection site in layers 5–6 (~500–600 μm deep), this angle would cause significant displacement along the horizontal axis (approximately 500–600 μm). Therefore, we chose the location of entry carefully, making sure that despite the horizontal displacement, the pipette would still remain within V1 once it reaches the desired depth.

#### 2.4.3 Pipette insertion into the brain

Once the entry location was in focus, the 40× objective lens was moved up by 2 mm to ensure the entry of the pipette. The pipette was mounted on a micromanipulator (MPC-365, Sutter Instrument) at an angle of 30° and the pipette tip containing the dye mixture was brought in focus under the objective lens using epi-fluorescence. Then the pipette was slowly lowered vertically using the manipulator while the objective lens was simultaneously lowered to keep the tip in focus, until the tip arrived at the pre-determined entry point on the brain surface. Two to three drops of agarose (2%, dissolved in artificial cerebrospinal fluid) were applied if brain pulsations were visible.

#### 2.4.4 Visualized travel to the injection site and dye injection

Once the tip was at the brain surface, imaging was switched to multiphoton mode and the two-photon excitation fluorescence from Alexa 633 at the pipette tip was visualized using 1300 nm excitation from the Spirit-NOPA. Under continuous visual guidance, the pipette was slowly advanced using the diagonal axis of the Sutter micromanipulator, until the pipette tip reached 500 μm below the brain surface. To prevent the pipette tip from clogging, occasional small pressure puffs (~10 psi) were applied as the pipette traversed through the brain. At the injection location, the dye mixture was pressure-ejected into the extracellular space (3–5 pulses, 30 s pulse duration, 10–15 psi per pulse) by using a Picospritzer (Parker Hannifin). Upon completion of injection, the pipette was withdrawn slowly, and a period of approximately one hour was allowed for the dye-loading of cells to complete. The craniotomy was sealed with agarose (1.5–2%, dissolved in artificial cerebrospinal fluid) and a 5 mm glass coverslip (World Precision Instruments) was placed over it. The mouse head was tilted back to horizontal position and the Olympus 25× objective lens was positioned over the labelled area before imaging commenced.

### 2.5 Visual stimulation

Drifting square-wave grating stimuli (100% contrast, 1.5 Hz temporal frequency and 0.025–0.033 cycles/degree) were presented on a 17-inch LCD monitor placed 15 cm from the eye. The stimuli were presented at 8 directions of motion in 45° steps. All our images were collected at 512 × 512 pixels. The sizes of the imaged fields of view (FOV) ranged between 265 × 265 and 495 × 495 μm, and the integration time per pixel was set to 3.2 μs. Under these imaging conditions, the frame periods ranged between 1.11 and 1.25 seconds. During functional imaging, drifting gratings were displayed for 5 imaging frames, interleaved with 10 frames of blank stimulus (equiluminant uniform grey), except for one experiment in which the blank stimulus was displayed for 7 frames. Thus, grating stimuli were displayed for approximately 6 seconds with 12 or 8 seconds of blank. Each condition was repeated at least 4 times.

### 2.6 Data analysis

All images were analyzed with custom code written in Matlab (Mathworks). Neurons were automatically masked by morphological criteria and validated manually^4, 6–7^. Fluorescence time courses of each neuron F(t) were calculated by averaging over the pixels within the mask. Then the response ΔF/F were computed as (F_1_−F_0_)/ F_0_, where F_1_ was the average fluorescence across the entire stimulus window and F_0_ was the average fluorescence during the last approximately 4 seconds from the blank interval. Responsive neurons were defined by ANOVA across 8 directions over multiple trials (*p* < 0.05).

To compute the direction selectivity, we first fit a double von Mises curve to the neural response:

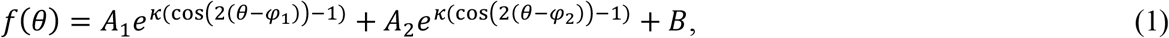

where *A*_1_ is the amplitude at the preferred direction, *A*_2_ is the amplitude of the second peak, *κ* represents a width parameter, *θ* is the orientation angle, *φ*_1_ is the preferred direction, *φ*_2_ is constrained to be *φ*_1_ +180°, and *B* is the baseline^25^. The response to the preferred direction of motion (*R*_*pref*_) and the opposite (null) direction of motion (*R*_*null*_) were estimated from the fitted curve.

The Direction Selectivity Index (DSI) was calculated as,

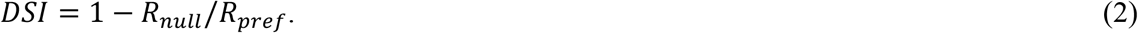

To determine Orientation Tuning Bandwidth (BW), responses to the two directions of motion for the same orientation were first averaged. Then, these responses were fit with single von Mises function (Equation 3 below).

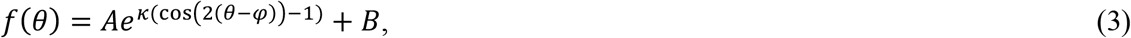

where *A* is the peak amplitude and *φ* is the preferred orientation. The other parameters are as in equation 1. BW was then calculated as the half-width at half-height of the fitted curve.

## 3. Results

We developed a non-genetic strategy for localizing calcium indicators in deeper cortical layers, to be subsequently utilized for recording neuronal activity via three-photon imaging. Briefly, we first mapped out the three-photon brightness spectra of the relevant fluorophores used in the experiments. Second, we optimized the fabrication of glass micropipettes with narrow profiles of the shank (see Methods). Then we filled these pipettes with OGB-1 AM and Alexa 633, and pressure-injected these dyes at 500–600 μm depths under multiphoton visualization. Finally, following bulk-loading of the neurons with OGB-1 AM, we performed three-photon fluorescence imaging of calcium signals at 1300 nm excitation and recorded visually-evoked responses.

Before we could use OGB-1 AM and other fluorophores for three-photon imaging, we needed to determine the best three-photon excitation wavelengths for these fluorophores. Theoretically, for any fluorophore, the peak excitation wavelengths for two- and three-photon absorption should be located at two and three times the one-photon absorption peak, respectively. However, two-photon absorption peaks for most fluorophores deviate from double the one-photon peaks^26–28^. This deviation may be true for three-photon absorption as well, thus necessitating actual measurement of the three-photon brightness spectra for relevant fluorophores to be used in experiments. Therefore, we mapped out the brightness spectra of a set of green and red fluorophores commonly used in imaging experiments, including the ones used in experiments described here (red fluorophores: Alexa 633, Texas Red Dextran, Sulforhodamine 101, mCherry, mRaspberry, tdTomato; green fluorophores: Oregon Green 488 BAPTA-1 hexapotassium salt, Fluorescein dextran). Briefly, we filled glass micropipettes with aqueous solutions of the dyes, imaged them under the microscope at varying excitation wavelengths and measured the fluorescence brightness at each wavelength (see Methods). For comparison, we measured the two-photon brightness spectra of the same dyes as well. The two-photon brightness spectra (Figure 2) revealed peaks that matched reported action cross-sections of the dyes^26–28^, validating that our method was sensitive in capturing the strong excitation wavelengths. The three-photon excitation peaks for the green fluorophores, including OGB-1 AM, were located around 1300 nm (Figure 2, wavelength range 1250–1350 nm). The red fluorophores revealed two broad peaks in their three-photon brightness spectra (Figure 2, wavelength range 1500–1800 nm). The precise location of the peaks varied slightly among the red fluorophores we tested. Moreover, while the two-photon spectra of all fluorophores revealed characteristic sharp peaks, the brightness peaks in the three-photon spectra were much broader. These broad peaks likely resulted from the fact that the excitation beam for three-photon excitation had much broader bandwidth (see Supplementary Fig. S1). From these spectra, we chose to use 1300 nm and 1600 nm as the optimal excitation wavelengths for three-photon imaging of OGB-1 AM and Texas Red Dextran, respectively. These wavelengths were also suitable because the absorption coefficient of water have local minima at 1300 and 1600 nm^29–30^, thereby causing less attenuation of the excitation beam due to water absorption.

**Figure 2.**
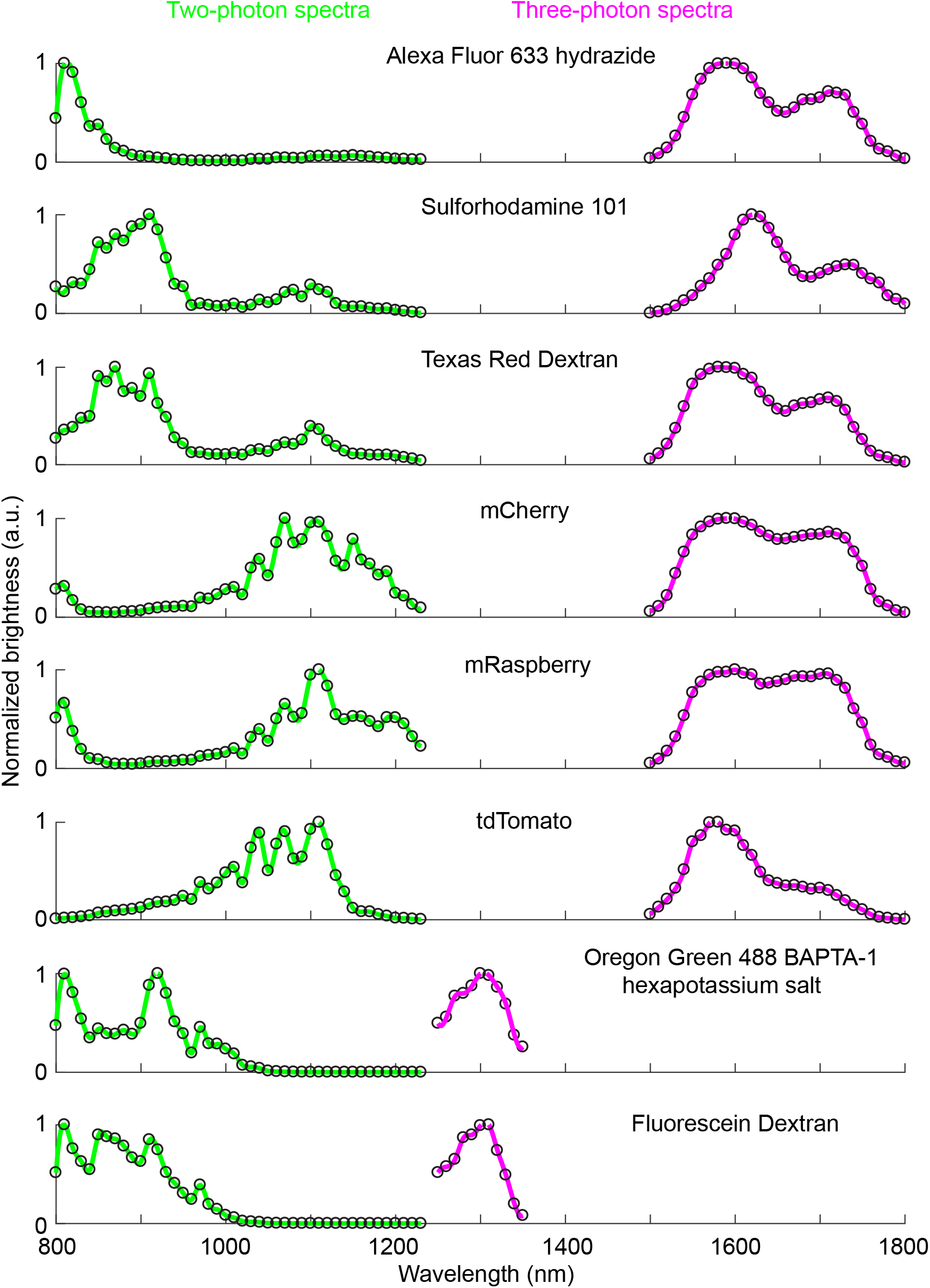
Two- and three-photon fluorescence brightness spectra of selected green and red fluorophores. Normalized two-photon (green curves) and three-photon (magenta curves) fluorescence brightness values are plotted against different excitation wavelengths. Open circles: fluorescent brightness values measured at excitation wavelengths 10 nm apart; continuous lines: individual points 10 nm apart interpolated at 2 nm steps. The measurements of each fluorophore were repeated at least twice and similar results were obtained.

Superficial injections of OGB-1 AM in layers 2/3 of mice and other species have been routinely carried out using glass micropipettes with 2–3 μm tip diameter^3–4^. In order to drive the pipettes down deeper into the cortex, it was important to fabricate pipettes with narrower shanks so that the damage to the superficial layers of cortex could be minimized. Therefore, we fabricated glass micropipettes with small tip diameters (2–3 μm) and slowly-tapering shanks (110 μm diameter at 1.4 mm behind the tip) that could be inserted deeper into the brain with limited damage to the tissue. With impedances ranging 13– 15 MΩ, these pipettes could be used to inject the dyes without clogging at the tips.

Injecting OGB-1 AM in layers 5–6 of the neocortex would require the pipette to travel through the brain significantly farther than what is required for conventional layer 2/3 injections. It is important to avoid piercing cells and blood vessels during the long travel to the injection depths, because this could lead to clogging of the tips and tissue damage. Therefore, we carried out the process of pipette insertion, travel and dye injection under continuous visualization of the pipette tip, which was achieved by mixing a red fluorescent dye (Alexa 633) with OGB-1 AM in the pipette and visualizing it via multiphoton imaging through a 40× objective (for details see Figure 3 and Methods). The mouse head was tilted to facilitate insertion of the pipette under the objective, and then a target location for pipette entry on the brain surface was chosen via bright-field imaging through the objective (Figure 3a). The objective was then retracted up by 2 mm, the pipette was advanced under the objective and the epi-fluorescence from Alexa 633 at the tip was brought into focus (Figure 3b). By simultaneously moving the pipette and the objective downward, the tip was then brought to the entry location on the brain surface (Figure 3c). At this point imaging modality was switched to multi-photon mode by using 1300 nm excitation, which generated strong two-photon excitation fluorescence from the Alexa 633 at the pipette tip. The pipette was then slowly advanced through the brain while maintaining the tip in focus, until the desired depth of injection was reached (Figure 3d). The dye mixture was then pressure-injected into the brain by applying air pulses from a Picospritzer. Following completion of the injection, the pipette was slowly retracted from the tissue. The cells were allowed to load with the dye for a period of 60 minutes, at the end of which the head was tilted back to horizontal orientation and multiphoton imaging of labelled cells commenced using a 25× objective (Figure 3e). Finally, in a subset of experiments, we also injected Texas Red Dextran into the bloodstream to image cortical blood vessels.

**Figure 3.**
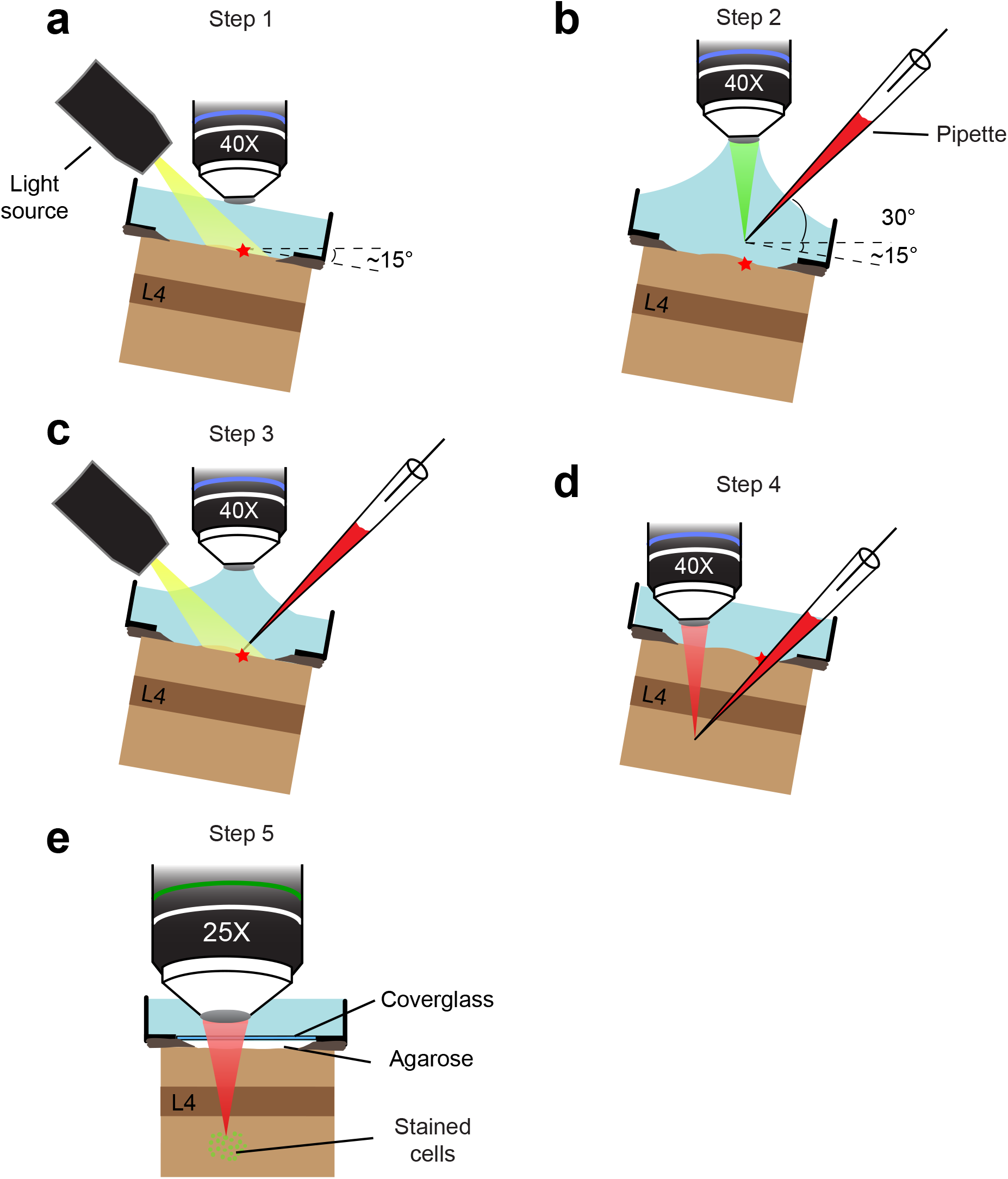
Schematic of the OGB-1 AM injection method. **a**, Step 1. The pipette entry point on the brain surface (red star) is determined by stereotaxic coordinates and anatomical landmarks. The surface vessels are visualized with a 40× objective under bright-field illumination. **b**, Step 2. The pipette tip is positioned directly above the entry point on the brain surface using green epi-fluorescent light after the objective is raised by 2 mm. **c**, Step 3. The pipette and the objective are lowered vertically until the brain surface comes into focus. **d**, Step 4. The pipette is driven down diagonally until its tip reaches the desired depth, here into cortical layer 5–6 located below layer 4 (L4). The tip is visualized throughout the travel via two-photon fluorescence of Alexa 633 under 1300 nm excitation. **e**, Step 5. After dye injection and labelling of cells, the craniotomy is sealed with agarose and a 5 mm glass coverslip. The head is tilted back to horizontal orientation and a 25× objective lens is positioned to perform three-photon imaging using 1300 nm excitation.

Various cortical compartments such as labelled cell bodies, blood vessels and fiber tracts were visualized using three imaging modalities. Three-photon imaging of OGB-1 AM fluorescence at 1300 nm excitation was used to visualize the activity of visual cortical neurons, and three-photon imaging of Texas Red Dextran fluorescence at 1600 nm excitation was used to visualize the entire vascular network. In addition, we used the label-free Third Harmonic Generation (THG) imaging at 1600 nm excitation to visualize blood vessels and a subset of the myelinated axons in the cortex. THG with 1600 nm excitation generated photon emission at 533 nm, which was detected in the green emission channel of the microscope (see Methods). We imaged an anatomical *z*-stack (n = 2 mice) of the entire cortical column using these three modalities and by varying power exponentially with imaging depth. The OGB-1 AM labelled a field of cells spanning ~300–400 μm in diameter localized to cortical layers 5–6 (Figure 4a, b, yellow). The vascular network could be imaged through the entire cortical column (Figure 4a–c, magenta). Blood vessels and some myelinated axons generated strong THG signals and the ventral boundary of the cortex could be easily demarcated through the clear THG visualization of the white matter tract (Figure 4a-c, cyan). The strong fluorescence from the vascular Texas Red Dextran allowed visualization of blood vessels running through the white matter tract (Figure 4c) as well as below it (Figure 4a).

**Figure 4.**
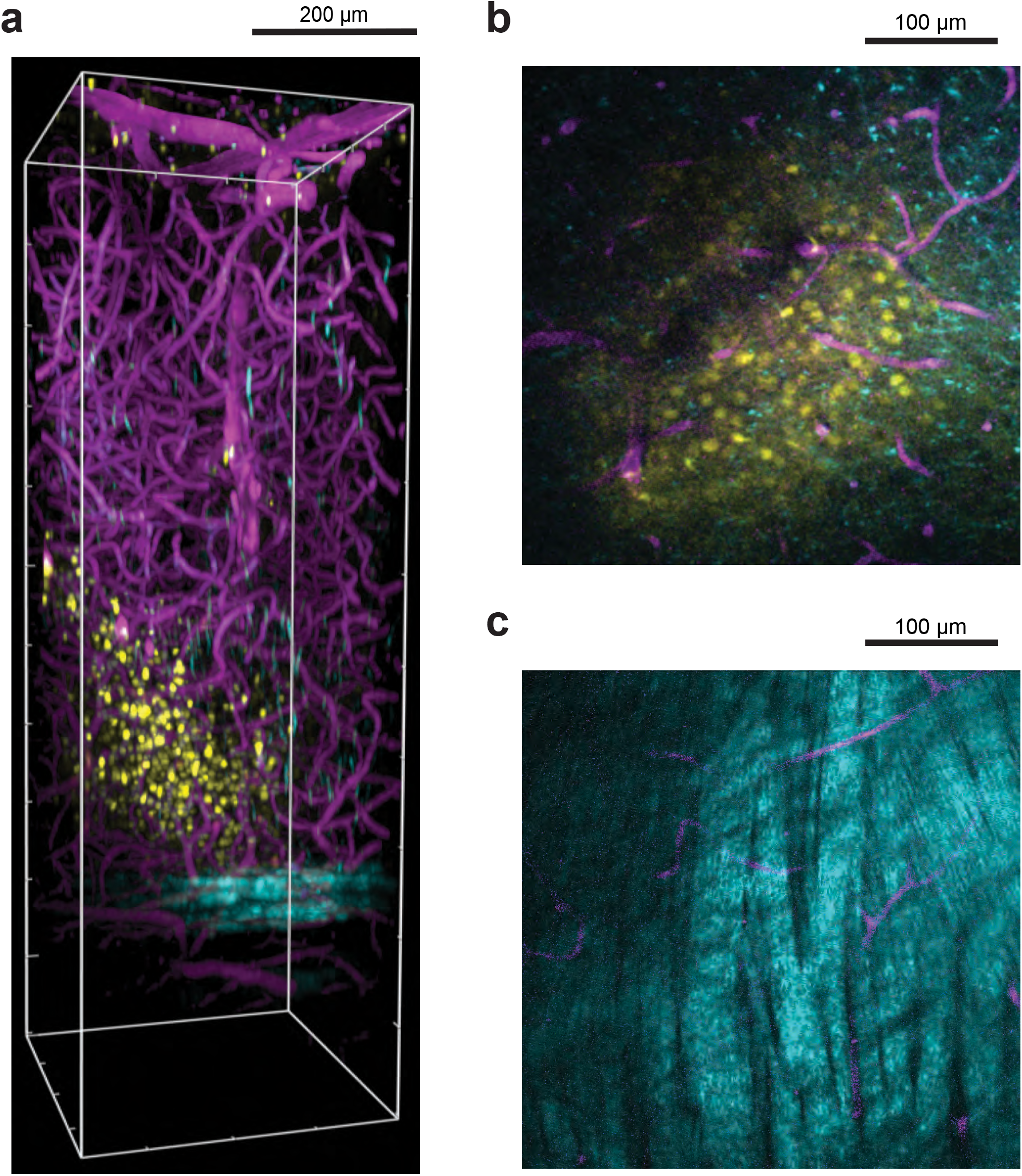
OGB-1 AM labelled cells deep in mouse visual cortex relative to blood vessels and white matter. **a**, Side view perspective of a three-dimensional reconstruction of the three-photon image volume depicting the field of OGB-labelled cells with respect to blood vessels and the white matter. OGB-1 AM labelled cells in layers 5–6 are shown in yellow, blood vessels labelled with Texas Red Dextran are shown in magenta and cortical white matter imaged with THG is shown in cyan. *x-y-z* volume dimensions: 375 μm × 375 μm × 1100 μm. **b**, Single *z* plane three-photon image at 750 μm below brain surface showing labelled cells and blood vessels. **c**, Single *z* plane three-photon image in the white matter at 900 μm below brain surface. Myelinated axons in the white matter have a banded structure and show a strong THG signal. The brightness and contrast were adjusted in **a**-**c** in order to highlight primarily either the labelled neurons (**a**, **b**) or the white matter tract (**c**).

To test the feasibility of recording neuronal responses using three-photon imaging of OGB-1 AM fluorescence, we presented drifting square-wave gratings in the contralateral monocular visual field of the mouse, while imaging the OGB-1 AM labelled cell fields. Functional imaging was carried out between the depths of 550–650 μm (n = 3 mice). Using 40–60 mW average power on the brain surface, we routinely recorded visual responses that were modulated by the grating orientation and direction of motion (Figure 5). Recent work has shown that for three-photon imaging at ~600 μm depth with 1300 nm excitation, heating-induced tissue damage starts between 100-150 mW average power^31^. Our use of 40–60 mW average power for OGB-1 AM imaging falls within those safety limits.

**Figure 5.**
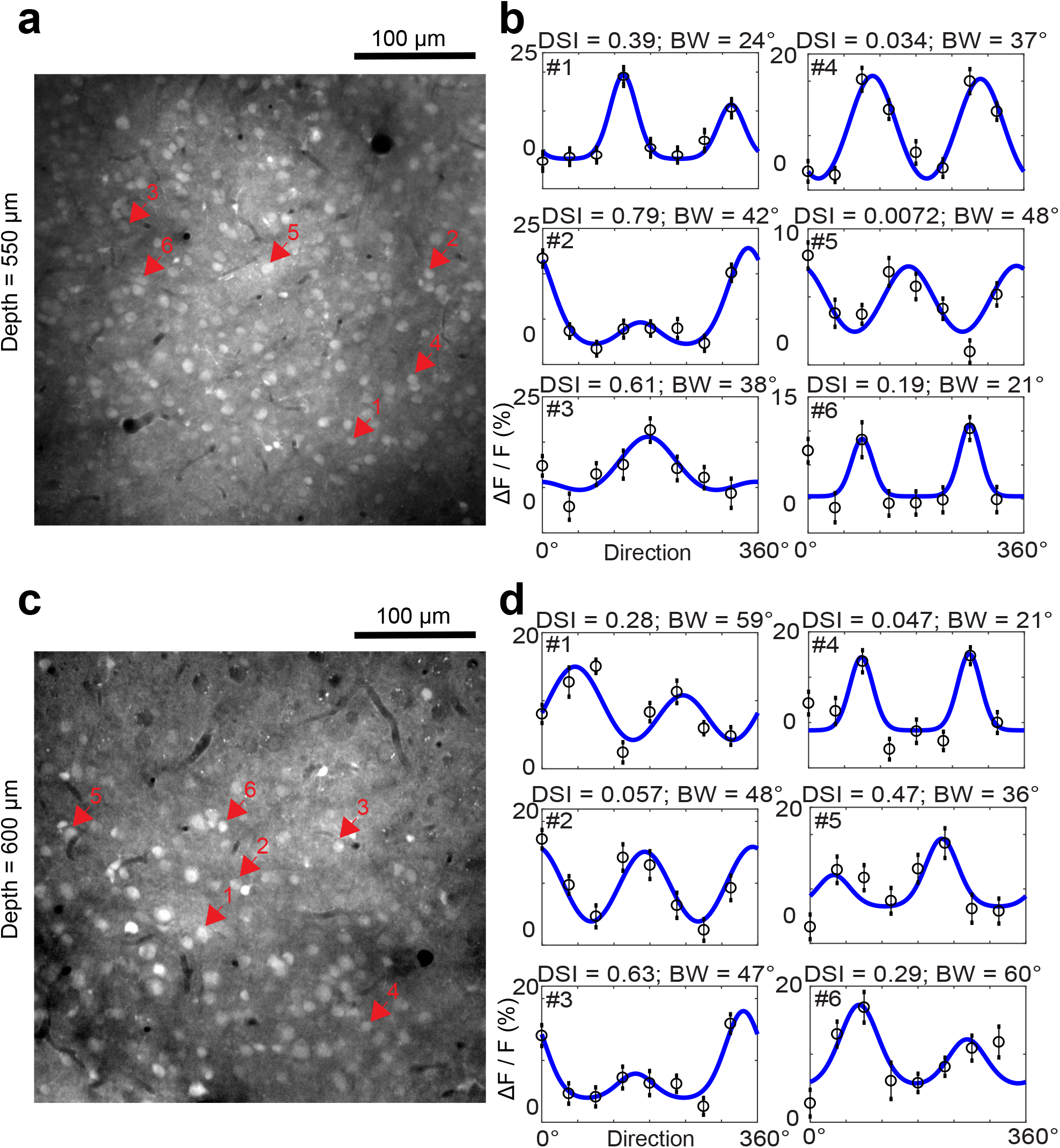
Visual responses of OGB-1 AM labelled neurons in layer 5–6 imaged by three-photon microscopy. **a**, Anatomical image at cortical depth 550 μm below the pial surface. **b**, Tuning curves of direction and orientation selectivity for six neurons (numbered 1 through 6, indicated by arrows in **a**). Direction Selectivity Index (DSI) and Orientation Tuning Bandwidth (BW, half-width at half-height) of each neuron are labelled accordingly. Open circles represent mean ΔF/F responses and vertical bars lines represent standard deviation; Blue curves are double Von Mises’ fits to the mean data. **c**, Anatomical image at cortical depth 600 μm below the pial surface. **d**, Tuning curves of direction and orientation selectivity for six neurons at 600 μm below pia (numbered 1 through 6, indicated by arrows in **c**).

## 4. Discussion

Our goal was to extend the use of calcium indicators in three-photon imaging to include bulk-loaded synthetic chemical dyes that could complement the use of GECIs, where applicable. We standardized the bulk-loading of layer 5–6 neurons in the mouse visual cortex with OGB-1 AM dye and subsequently imaged visual responses from labelled neurons using three-photon microscopy at 1300 nm excitation. Our results show that OGB-1 AM can be used for three-photon functional imaging of deeper cortical layers.

The choice of optimal excitation wavelength is of critical importance in three-photon microscopy, and is not driven by the consideration of the absorption cross-section of the fluorophore alone. The near infra-red (NIR) wavelengths of light suitable for three-photon excitation are absorbed strongly by water, which attenuates the excitation beam intensity and leads to significant heating of the brain tissue. The absorption spectrum of water in the NIR range contains several peaks and valleys, revealing two narrow regions centered around 1300 nm and 1700 nm where excitation attenuation is minimum^16, 29–30, 32^. The ultimate choice of excitation wavelength for a particular fluorophore is based on a combination of this attenuation window and the absorption cross-section of the fluorophore. Ideally, the optimal wavelength would be one where the fluorophore absorption is high (peak in absorption cross-section spectrum) and water absorption is low (valley in water absorption spectrum). However, for some fluorophores, if the peak in fluorophore absorption does not coincide with the trough in water absorption, depending on the relative amplitudes of the fluorophore absorption spectrum a decision has to be made regarding what wavelength to use in three-photon imaging.

Therefore, we first mapped out the optimal three-photon excitation wavelengths for the fluorophores used in our experiment (OGB-1 AM, Alexa 633 and Texas Red Dextran) and some other fluorescent dyes and proteins routinely used in multiphoton imaging. Conventionally, the intrinsic fluorescent properties (absorption cross-section) of fluorophores are quantified through spectroscopic methods, but the results from study to study varies tremendously (reviewed by Drobizhev and colleagues^33^). For the practical utilization of fluorophores in a multiphoton imaging context, imaged fluorescence brightness under varying wavelengths of excitation reveals information sufficient for optimizing the excitation wavelength used for imaging experiments. For example, the brightness spectra shown in Figure 2 indicated that the best excitation wavelength to be used for a specific fluorophore under the optical parameters of our imaging system design, which includes the lasers, table optics and microscope. The peak brightness wavelengths for the same fluorophore may be different in other imaging systems which may be configured differently. The fluorescence brightness imaged inside a glass micropipette or cuvette is not solely dependent on the absorption cross-section of the fluorophore, instead it depends on various other imaging parameters^34^. For example, the fluorescence brightness in three-photon excitation is highly sensitive to the pulse duration of the excitation laser beam. But the pulse duration of an excitation beam under the objective lens could vary across different wavelengths and from one imaging system to another, depending on the dispersion properties of the optics used. As a consequence, the wavelength that produces the peak brightness could also differ between different imaging systems. Therefore, the brightness spectra we report are operational guides for one particular imaging system, they should not be confused with universal action cross-section plots, and such spectra need to be determined independently for each imaging system. For green fluorophores such as OGB-1 AM in our setup, the choice of wavelength was simple as there was only one peak in the brightness spectrum around 1300 nm, which also coincides well with the local minima in water absorption. For red fluorophores, we saw two broad peaks centered around 1600 and 1700 nm, the relative amplitude of the peaks varying between fluorophores. For Texas Red Dextran, the bigger peak fell near 1600 nm, where the effective attenuation length of the mouse cortex happens to be slightly shorter than at 1700 nm^32^, implying that there is slightly more attenuation at 1600 nm. However, the brightness peak at 1600 nm was almost twice in amplitude compared to 1700 nm. Therefore, we chose 1600 nm as optimal excitation wavelength for Texas Red Dextran.

We succeeded in restricting the OGB-1 AM loading to cells within a diameter of 300-400 μm from the injection site to cortical layers 5–6. As a result, the superficial layers of cortex above the loaded cell field contained no labelled cells and thus contributed little or no fluorescence. This situation differs from the previously reported studies using virally expressed GECIs^19^, where neurons in layers 2/3 were also labelled with GCaMP6. Despite strong non-linear confinement of the excitation beam within a very narrow focal volume in three-photon excitation, superficial fluorophores in layers 2/3 would nevertheless contribute some low levels of fluorescence, thereby reducing SBR and attenuating the excitation beam for deeper cortical layers. The localized labelling achieved by our method, in contrast, ensured three-photon excitation fluorescence from only the labelled cell field localized at the desired depth. The consequent marginal improvements in SBR may prove critical when performing three-photon imaging even deeper in the brain, or in other types of tissues which may have even shorter attenuation lengths.

Functional imaging of calcium signals through OGB-1 AM fluorescence in deeper cortical layers, as described here, overcome the challenges of imaging with some GECIs. The AAV-based viral approaches typically use synapsin or CaMKII promoters to drive expression of the GECIs in mammalian neurons. In the neocortex of most mammals tested so far, these promoters drive strong expression in layers 1, 2, 3, 5 and 6, but in layer 4 only a small subset of neurons express the exogenous protein (see Anderman et al., 2013 ^35^ and Supplementary Fig. S2 for mouse and cat neocortex). Because two-photon imaging does not provide access to depths at which cortical layer 4 is typically located in non-rodent species, the lack of reliable expression in layer 4 has not been a major impediment to two-photon imaging experiments thus far. But now that three-photon imaging has made it possible to carry out high-resolution optical imaging at depths of 1 mm or beyond, layer 4 neurons in non-rodent species such as cats, ferrets or macaques (typically spanning 750–1200 μm from the surface) are within imaging access, provided one can label these neurons with a calcium indicator. In this regard, the lack of virally-driven expression in layer 4 poses a challenge for using GECIs to record layer 4 neuronal activity through three-photon imaging. The method described here, involving bulk-loading of a synthetic calcium indicator through localized injection, could provide a viable alternative approach to label neurons in and carry out functional three-photon imaging from layer 4 neurons in ferrets, cats and monkeys.

## Acknowledgements

We thank Austin Leikvoll for histology, technical assistance during mouse surgeries and comments on the manuscript. This research was supported by grants from the NIH (R01 MH111447) and NSF (1707287).

## Author contributions

CJL, AR, DMF and PK designed experiments, CJL, AR and DMF optimized the table optics, CJL, AR, AAS and DMF collected data, CJL, AR, AAS and DMF performed data analysis, CJL, AR and PK wrote the manuscript. All authors reviewed and approved the final submitted manuscript.

## References

1. W. Denk, J. H. Strickler, and W. W. Webb, “Two-photon laser scanning fluorescence microscopy,” Science 248(4951), 73–76 (1990).

2. E. Brustein et al., ““In vivo” monitoring of neuronal network activity in zebrafish by two-photon Ca(2+) imaging,” Pflugers Arch 446(6), 766–773 (2003).

3. C. Stosiek et al., “In vivo two-photon calcium imaging of neuronal networks,” Proc Natl Acad Sci USA 100(12), 7319–7324 (2003).

4. K. Ohki et al., “Functional imaging with cellular resolution reveals precise micro-architecture in visual cortex,” Nature 433(7026), 597–603 (2005).

5. V. Bonin et al., “Local diversity and fine-scale organization of receptive fields in mouse visual cortex,” J Neurosci 31(50), 18506–18521 (2011).

6. K. Ohki et al., “Highly ordered arrangement of single neurons in orientation pinwheels,” Nature 442(7105), 925–928 (2006).

7. P. Kara, and J. D. Boyd, “A micro-architecture for binocular disparity and ocular dominance in visual cortex,” Nature 458(7238), 627–631 (2009).

8. P. O’Herron et al., “Neural correlates of single-vessel haemodynamic responses in vivo,” Nature 534(7607), 378–382 (2016).

9. Y. Li et al., “Experience with moving visual stimuli drives the early development of cortical direction selectivity,” Nature 456(7224), 952–956 (2008).

10. G. B. Smith et al., “The development of cortical circuits for motion discrimination,” Nat Neurosci 18(2), 252–261 (2015).

11. I. Nauhaus et al., “Orthogonal micro-organization of orientation and spatial frequency in primate primary visual cortex,” Nat Neurosci 15(12), 1683–1690 (2012).

12. M. Li et al., “Long-Term Two-Photon Imaging in Awake Macaque Monkey,” Neuron 93(5), 1049–1057 (2017).

13. M. Oheim et al., “Two-photon microscopy in brain tissue: parameters influencing the imaging depth,” J Neurosci Methods 111(1), 29–37 (2001).

14. P. Theer, and W. Denk, “On the fundamental imaging-depth limit in two-photon microscopy,” J Opt Soc Am A Opt Image Sci Vis 23(12), 3139–3149 (2006).

15. K. Takasaki, R. Abbasi-Asl, and J. Waters, “Superficial Bound of the Depth Limit of Two-Photon Imaging in Mouse Brain,” eNeuro 7, ENEURO.0255–0219.2019 (2020).

16. N. G. Horton et al., “In vivo three-photon microscopy of subcortical structures within an intact mouse brain,” Nat Photonics 7(3), 205–209 (2013).

17. S. W. Hell et al., “Three-photon excitation in fluorescence microscopy,” J Biomed Opt 1(1), 71–74 (1996).

18. C. Xu et al., “Multiphoton fluorescence excitation: new spectral windows for biological nonlinear microscopy,” Proc Natl Acad Sci USA 93(20), 10763–10768 (1996).

19. M. Yildirim et al., “Functional imaging of visual cortical layers and subplate in awake mice with optimized three-photon microscopy,” Nat Commun 10(1), 177 (2019).

20. K. T. Takasaki, D. Tsyboulski, and J. Waters, “Dual-plane 3-photon microscopy with remote focusing,” Biomed Opt Express 10(11), 5585–5599 (2019).

21. D. G. Ouzounov et al., “In vivo three-photon imaging of activity of GCaMP6-labeled neurons deep in intact mouse brain,” Nat Methods 14(4), 388–390 (2017).

22. T. Wang et al., “Three-photon imaging of mouse brain structure and function through the intact skull,” Nat Methods 15(10), 789–792 (2018).

23. D. E. Wilson et al., “GABAergic Neurons in Ferret Visual Cortex Participate in Functionally Specific Networks,” Neuron 93(5), 1058–1065 (2017).

24. P. O’Herron et al., “Targeted labeling of neurons in a specific functional micro-domain of the neocortex by combining intrinsic signal and two-photon imaging,” J Vis Exp 70, e50025 (2012).

25. N. V. Swindale, “Orientation tuning curves: empirical description and estimation of parameters,” Biol Cybern 78(1), 45–56 (1998).

26. K. Svoboda, and R. Yasuda, “Principles of two-photon excitation microscopy and its applications to neuroscience,” Neuron 50(6), 823–839 (2006).

27. M. Drobizhev et al., “Absolute two-photon absorption spectra and two-photon brightness of orange and red fluorescent proteins,” J Phys Chem B 113(4), 855–859 (2009).

28. J. Mutze et al., “Excitation spectra and brightness optimization of two-photon excited probes,” Biophys J 102(4), 934–944 (2012).

29. L. Kou, D. Labrie, and P. Chylek, “Refractive indices of water and ice in the 0.65- to 2.5-microm spectral range,” Appl Opt 32(19), 3531–3540 (1993).

30. D. Kobat et al., “Deep tissue multiphoton microscopy using longer wavelength excitation,” Opt Express 17(16), 13354–13364 (2009).

31. T. Wang et al., “Quantitative analysis of 1300-nm three-photon calcium imaging in the mouse brain,” eLife 9, e53205 (2020).

32. M. Wang et al., “Comparing the effective attenuation lengths for long wavelength in vivo imaging of the mouse brain,” Biomed Opt Express 9(8), 3534–3543 (2018).

33. M. Drobizhev et al., “Two-photon absorption properties of fluorescent proteins,” Nat Methods 8(5), 393–399 (2011).

34. C. Xu, and W. W. Webb, “Measurement of two-photon excitation cross sections of molecular fluorophores with data from 690 to 1050 nm,” J Opt Soc Am B 13(3), 481–491 (1996).

35. M. L. Andermann et al., “Chronic cellular imaging of entire cortical columns in awake mice using microprisms,” Neuron 80(4), 900–913 (2013).

