## Supplemental material for "Three-photon imaging of synthetic dyes in deep layers of the neocortex"

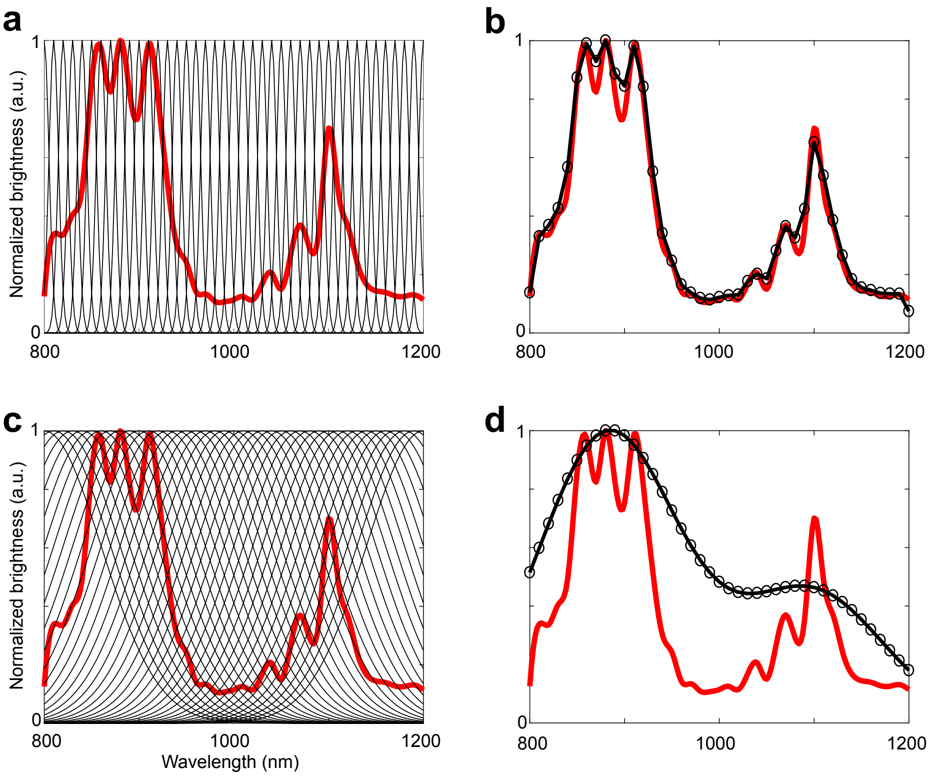


**Supplementary Figure S1**. Simulation results showing that the wider bandwidth of the three-photon excitation beam may cause the broad peaks in the three-photon brightness spectra.

Two-photon excitation spectra for fluorophores typically display sharp peaks (see Figure 1 green curves). In contrast, we found broad peaks in the three-photon spectra of most fluorophores we tested (see Figure 1 magenta curves). One potential explanation lies in the difference in the bandwidth of the excitation beams that exists between two- and three-photon excitation sources. While the bandwidth for two-photon excitation beams (800-1200 nm central wavelengths) varies between 5-20 nm, the bandwidth for three-photon excitation beams (1250-1800 nm central wavelengths) varies between 60-80 nm. Probing a fluorophore's excitation spectrum with broad-bandwidth beams will result in down-sampling the spectrum and yield broader peaks. To test this idea, we ran simulations on a hypothetical underlying spectrum with sharp peaks using narrow-bandwidth (5 nm) or broad-bandwidth (60 nm) excitation beams, and compared the resulting output spectra.

a) Experimentally measured 2-photon spectrum of Texas Red Dextran versus simulated narrow-bandwidth beams to probe the spectrum. The red line represents the two-photon brightness spectrum of Texas red dextran, used in the simulation as a generic underlying "true" spectrum being probed during spectrum mapping by exposing it to various excitation wavelengths. The black lines are Gaussian functions with 5 nm bandwidth, centered on 800 to 1200 nm wavelengths at 10 nm steps, representing the beams being used for probing the fluorophore.

b) Outcome of probing the Texas Red spectrum with narrow-bandwidth excitation beams. Red line: same as in panel **a**; black line: the results of convolving the Texas red two-photon spectrum and the Gaussian functions in panel **a**. The black line in panel **b** simulates the brightness spectrum that will result from probing the "true" spectrum with excitation beams at 5 nm bandwidth. The output spectrum closely follows the contours of the original spectrum.

c–d) Experimentally measured 2-photon spectrum of Texas Red Dextran probed with wide-bandwidth beams. Panels **c** and **d** are similar to panels **a** and **b**, respectively, except that the excitation beams in panel **c** are with 60 nm bandwidth. The black line in panel **d** simulates the brightness spectrum resulting from probing the "true" spectrum with excitation beams with 60 nm bandwidth. The output spectrum in this case reveals slowly-varying broad peaks whereas the original spectrum displays sharp peaks.

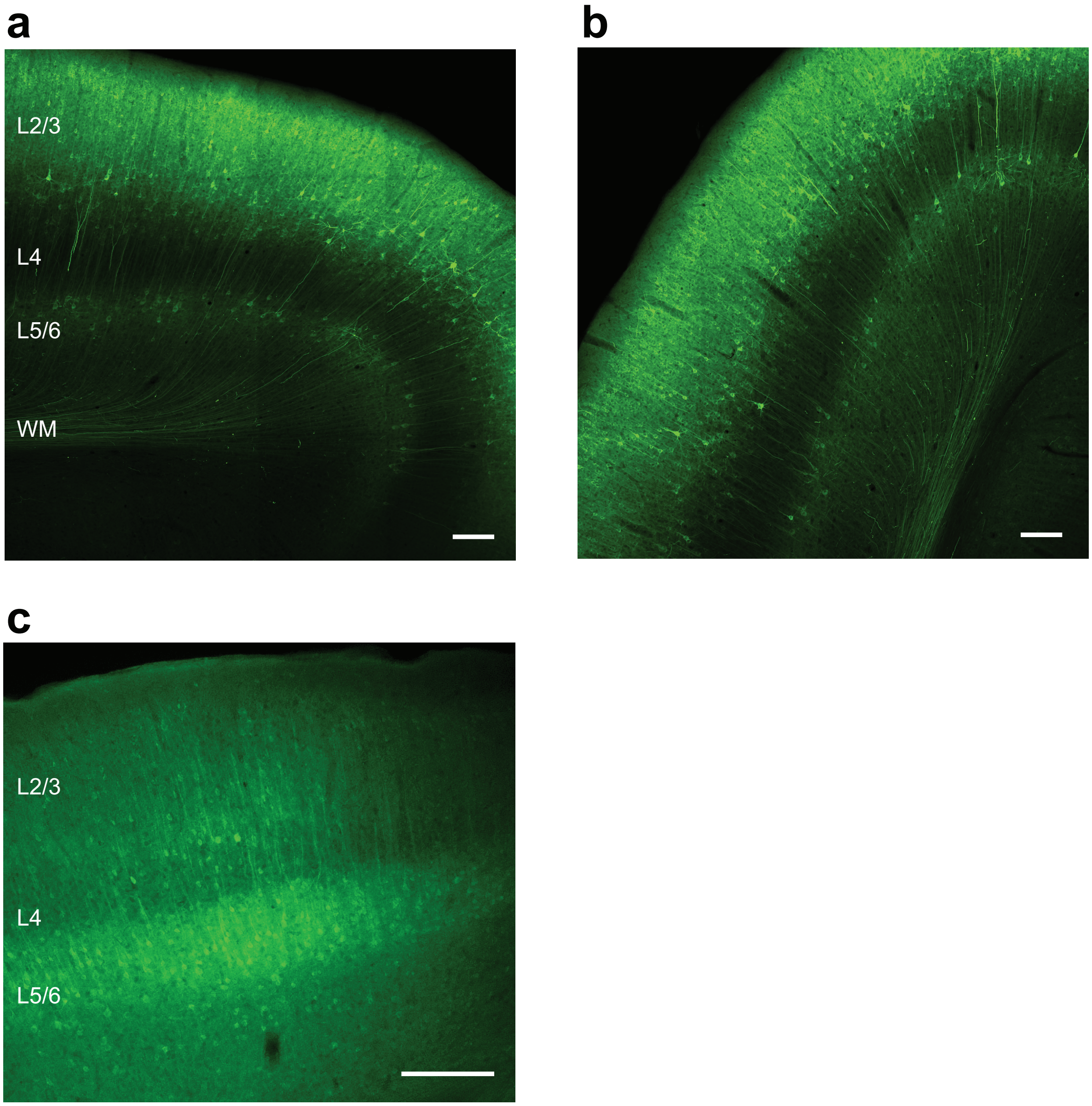


**Supplementary Figure S2**. Viral transfection-mediated GCaMP6s expression is weak in layer 4 of the cat and mouse visual cortex.

a-b) Sample images of GCaMP6s expression in two brain slices from the primary visual cortex (V1) area of one cat. The slices were collected from a cat brain used in a previous study (*O'Herron et. al., Nature 534, 378-382, 2016*). GCaMP6s expression was driven under the synapsin promoter packaged inside an adeno-associated virus (AAV). In both slices, strong GCaMP6s expression can be detected in many neurons and the neuropil in layers 2/3 and 5/6. In contrast, layer 4 contains very few weakly-labelled cells and the general intensity of GCaMP6s fluorescence in layer 4 was low. WM: white matter. Scale bars: 200 µm.

c) A brain slice from the mouse V1 showing GCaMP6s expression driven by the same viral vector as in **a** and **b**. Similar pattern of low GCaMP6s expression in layer 4 is observed in the mouse V1. Scale bar: 200 µm.

Paraformaldehyde-fixed brain slices (50 µm thick) were mounted on glass slides to image the native fluorescence from the GCaMP6s protein. The fluorescence was excited using 488 nm light and the emission was detected at 525 nm. The cat brain slices were imaged with a Nikon A1R confocal microscope and a Nikon Plan Apo 20× objective (0.75 NA). An approximately 2.5 mm × 2.5 mm area of each slice was tiled 4×4 and at each location a z-stack was collected covering the slice thickness at 2 µm steps. Nikon NIS Elements software was used to stitch together the individual images with 5% overlap, and a maximum intensity z projection applied to obtain the final images. The epi-fluorescence of the mouse brain slices were imaged with a Bruker microscope and an Olympus UPLFLN 4× objective (0.13 NA).
